# A functional analysis of the Drosophila gene *hindsight*: evidence for positive regulation of EGFR signaling

**DOI:** 10.1101/814137

**Authors:** Minhee Kim, Olivia Y. Du, Rachael J. Whitney, Ronit Wilk, Jack Hu, Henry M. Krause, Joshua Kavaler, Bruce H. Reed

**Affiliations:** Department of Biology, University of Waterloo, Waterloo, ON, Canada N2L 3G1.; Department of Molecular Genetics, The Terrence Donnelly Centre for Cellular and Biomolecular Research, University of Toronto, Toronto, ON, Canada, M5S 3E1; Department of Biology, Colby College, Waterville, ME, USA 04901

**Keywords:** Hindsight/RREB-1, EGFR signaling, MAPK, germ band retraction

## Abstract

We have investigated the relationship between the function of the gene *hindsight* (*hnt*), which is the Drosophila homolog of *Ras Responsive Element Binding protein-1* (*RREB-1*), and the EGFR signaling pathway. We report that *hnt* mutant embryos are defective in EGFR signaling dependent processes, namely chordotonal organ recruitment and oenocyte specification. We also show the temperature sensitive hypomorphic allele *hnt^pebbled^* is enhanced by the hypomorphic MAPK allele *rolled* (*rl^1^*). We find that *hnt* overexpression results in ectopic *DPax2* expression within the embryonic peripheral nervous system, and we show that this effect is EGFR-dependent. Finally, we show that the canonical U-shaped embryonic lethal phenotype of *hnt,* which is associated with premature degeneration of the extraembyonic amnioserosa and a failure in germ band retraction, is rescued by expression of several components of the EGFR signaling pathway (*sSpi*, *Ras85D^V12^*, *pnt^P1^*) as well as the caspase inhibitor *p35*. Based on this collection of corroborating evidence, we suggest that an overarching function of *hnt* involves the positive regulation of EGFR signaling.

## Introduction

The gene *hindsight* (*hnt*), also known as *pebbled* (*peb*), was first identified in mutagenesis screens for embryonic lethal mutations performed in the early 1980’s (WIESCHAUS *et al*. 1984). The embryonic lethal phenotype of *hnt* was categorized as “U-shaped”, reflecting a failure to undergo or complete germ band retraction. *hnt* has since been identified as the Drosophila homolog of mammalian *Ras Responsive Element Binding Protein −1* (*RREB-1*) (MELANI *et al*. 2008; MING *et al*. 2013), which strongly suggests a connection between *hnt* and the EGFR/Ras/MAPK signaling pathway (hereafter referred to as EGFR signaling). Interestingly, in Drosophila, *hnt* has been identified as a direct transcriptional target of the Notch signaling pathway (KREJCI *et al*. 2009; TERRIENTE-FELIX *et al*. 2013). Mammalian *RREB-1*, on the other hand, has not been linked with Notch signaling but functions downstream of Ras/MAPK signaling and may either activate or repress certain Ras target genes (LIU *et al*. 2009; KENT *et al*. 2014). *RREB-1* has also been implicated in a number of human pathologies, including pancreatic, prostate, thyroid, and colon cancer (THIAGALINGAM *et al*. 1996; MUKHOPADHYAY *et al*. 2007; KENT *et al*. 2013; FRANKLIN *et al*. 2014).

The *hnt* gene encodes a transcription factor composed of 1893 amino acids containing 14 C_2_H_2_-type Zinc-fingers (YIP *et al*. 1997). Based on genetic interaction studies, Hnt’s target genes are likely numerous and disparate with respect to function (WILK *et al*. 2004). Candidate direct target genes of Hnt identified using molecular methods include *hnt* itself, *nervy*, and *jitterbug* (MING *et al*. 2013; OLIVA *et al*. 2015). The *nervy* gene encodes a *Drosophila* homolog of the human proto-oncogene ETO/MTG8, while *jitterbug* encodes a conserved actin binding protein also known as *filamen*.

During development *hnt* is expressed in a broad range of tissues. In the embryo these include the amnioserosa (AS), anterior and posterior midgut primordia, the peripheral nervous system (PNS), the developing tracheal system, and the oenocytes (YIP *et al*. 1997; WILK *et al*. 2000; BRODU *et al*. 2004). During larval stages, in addition to the tracheal system, PNS, midgut, and oenocytes, *hnt* is expressed in the larval lymph gland, differentiated crystal cells, imaginal tracheoblasts, and the salivary glands of the third instar (PITSOULI AND PERRIMON 2010; MING *et al*. 2013; TERRIENTE-FELIX *et al*. 2013). In pupae, the sensory organ precursors (SOPs) of developing micro- and macrochaetae, as well as myoblasts, and all photoreceptor cells (R cells) of the developing retina express *hnt* (PICKUP *et al*. 2002; REEVES AND POSAKONY 2005; KREJCI *et al*. 2009; BUFFIN AND GHO 2010). In the adult, Hnt is expressed in the midgut (intestinal stem cells, enteroblasts, and enterocytes), developing egg chambers (follicle cells and the migratory border cells), spermathecae, and in mature neurons of the wing (SUN AND DENG 2007; MELANI *et al*. 2008; BAECHLER *et al*. 2015; SHEN AND SUN 2017; FARLEY *et al*. 2018).

While *hnt* is expressed in many different tissues, its expression within a given tissue can be dynamic. For example, in the adult intestinal stem cell lineage there is an increase of Hnt during enteroblast-to-enterocyte differentiation, but a decrease during enteroblast-to-enteroendocrine cell differentiation (BAECHLER *et al*. 2015). Hnt levels are particularly dynamic in the ovarian follicle cells, where Hnt is observed in stage 7-10A egg chambers as these cells initiate endoreduplication. A subset of follicle cells are subsequently devoid of Hnt through stages 10B to 13, and then display a strong increase in stage 14 egg chambers prior to follicle cell rupture and an ovulation-like event (DEADY *et al*. 2017).

There is a wealth of information regarding *hnt* mutant phenotypes and *hnt* expression, yet a general definition of Hnt function remains elusive. Given that Hnt is the Drosophila homolog of RREB-1, we present an examination of *hnt* mutant phenotypes as well as *hnt* overexpression with specific attention to EGFR signaling. With respect to loss-of function analysis, we report two new findings that link *hnt* and EGFR signaling: first, *hnt* mutant embryos are defective in the processes of chordotonal organ recruitment as well as oenocyte specification, both of which are EGFR signaling-dependent processes (MAKKI *et al*. 2014); and second, we show that the temperature sensitive *hnt* allele *hnt^pebbled^* (*hnt^peb^*), which is associated with defective cone cell specification in the pupal retina (PICKUP *et al*. 2009), is enhanced by the hypomorphic MAPK allele *rolled* (*rl^1^*). In terms of *hnt* overexpression, we first show ectopic *DPax2* expression in embryos overexpressing *hnt.* We show similar ectopic *DPax2* expression in embryos in which EGFR signaling is abnormally increased through global expression of the active EGFR ligand *secreted Spitz* (*sSpi*). We subsequently demonstrate that *Egfr* loss-of-function mutants abrogate ectopic *DPax2* expression in the context of *hnt* overexpression. Last, we show that the U-shaped phenotype of *hnt* mutants, which involves premature degeneration of the AS and a failure in the morphogenetic process of germ band retraction (GBR) - which is also a phenotype displayed by *Egfr* mutants (CLIFFORD AND SCHUPBACH 1992) - can be rescued by expression of components of the EGFR signaling pathway (*sSpi*, *Ras85D^V12^*, *pnt^P1^*) as well as the caspase inhibitor *p35*. Interestingly, expression of the *pnt^P2^* isoform, which (unlike the *pnt^P1^* isoform) requires activation by MAPK (O’NEILL *et al*. 1994; SHWARTZ *et al*. 2013), does not rescue *hnt* mutants. Given this collection of corroborating evidence, we suggest that a primary function of *hnt* involves the positive regulation of EGFR signaling.

## Materials and Methods

### Drosophila stocks

All cultures were raised on standard *Drosophila* medium at 25°C under a 12 hour light/dark cycle, unless otherwise indicated. The *hindsight* (*hnt*) alleles used were *hnt^XE81^, hnt^peb^* (YIP *et al*. 1997; WILK *et al*. 2004), and *hnt^NP7278ex1^* (this study). As previously described (YIP *et al*. 1997), *hnt^XE81^* is a strong hypomorphic embryonic lethal allele while *hnt^peb^* is a viable temperature sensitive hypomorphic allele associated with a rough eye phenotype at the restrictive temperature of 29° C. The *Egfr* mutant alleles used were *Egfr^1a15^* and *Egfr^f2^* as previously described (SHEN *et al*. 2013). The *rolled* (*rl^1^*) allele was provided by A. Hilliker. To drive ubiquitous expression throughout the early embryo we used *daGAL4* as previously described (REED *et al*. 2001). The *BO-GAL4* line was used to mark embryonic oenocytes (GUTIERREZ *et al*. 2007) and was provided by A. Gould. Overexpression of *hnt* used *UAS-GFP-hnt* as previously described (BAECHLER *et al*. 2015). The adherens junctions marker *Ubi-DEcadherin-GFP* was used to outline cell membranes as previously described (CORMIER *et al*. 2012). The reporter gene *DPax2^B1^GFP* was as previously described (JOHNSON *et al*. 2011). *UAS-sSpi* was obtained from N. Harden. *pebBAC^CH321-46J02^* was obtained from M. Freeman. All other transgenes used originated from stocks obtained from the Bloomington Drosophila Stock Center (*UAS-CD8-GFP*, *UAS-GFP^nls^*, *UAS-p35*, *UAS-Ras85D^V12^*, *UAS-pnt^P1^*, *UAS-pnt^P2^*)

### Construction of *DPax2-dsRed* reporter lines

The *DPax2^B1^dsRed* and *DPax2^B2^dsRed* reporter lines were generated by standard *P*-element transgenic methods (BACHMANN AND KNUST 2008) using the vector pRed H-Stinger (BAROLO *et al*. 2004) containing a previously described 3 KB *DPax2* enhancer (JOHNSON *et al*. 2011). Briefly, the 3 KB enhancer (position −3027 to +101 relative to the DPax2 transcription start site) was excised from the Bam HI sites of a DPax^B^-pBluescript KS + plasmid. The insert was then cloned into the Bam HI site of pRed H-Stinger.

### Crossing schemes for analysis of *DPax2^B2^dsRed* expression in *Egfr* mutants, and *DPax2^B1^GFP* expression in embryos with elevated EGFR signaling

In order to analyze *DPax2* reporter construct expression in different backgrounds, the *Ubi-DEcadherin-GFP* (on *second* chromosome) was recombined with *Egfr^1a15^*, *UAS-GFP-hnt* (on *second* chromosome) was recombined with *Egfr^f2^*, *daGAL4* (on *third* chromosome) was recombined with *DPax2^B2^dsRed*, and *daGAL4* (on *third* chromosome) was recombined with *DPax2^B1^GFP* creating the following stocks:

**Stock 1**: *dp^1a15^ Ubi-DEcadherin-GFP Egfr^1a15^*/ *CyO*

**Stock 2**: *UAS-GFP-hnt Egfr^f2^*/ *CyO*

**Stock 3**: *daGAL4 DPax2^B2^dsRed*

**Stock 4**: *daGAL4 DPax2^B1^GFP* / *TM6C*

To visualize *DPax2^B2^dsRed* expression in *Egfr^1a15^*/*Egfr^f2^* mutants, as well as *Egfr^f2^*/*+* heterozygotes, the following approach was used. Non-balancer male progeny of Stock 1 x Stock 3 (*dp^1a15^ Ubi-DE-cadherin Egfr^1a15^*/*+*; *daGAL4 DPax2^B2^dsRed*/*+*) were crossed to Stock 2. In embryos collected from this cross, *Egfr^1a15^*/*Egfr^f2^* mutants were recognized as embryos expressing *UAS-GFP-hnt*, *DPax2^B2^dsRed*, and *Ubi-DE-cadherin-GFP*, while *Egfr^f2^*/*+* heterozygotes also expressed *UAS-GFP-hnt* and *DPax2^B2^dsRed*, but lacked *Ubi-DE-cadherin-GFP*.

To visualize *DPax2^B1^GFP* expression in embryos with elevated EGFR signaling, Stock 4 was crossed to homozygous *UAS-sSpi*.

### Immunostaining and Imaging

Immunostaining of embryos was carried out as described (REED *et al*. 2001). The following primary antibodies were used at the indicated dilutions: mouse monoclonal anti-Hindsight (Hnt) 27B8 1G9 (1:25; from H. Lipshitz, University of Toronto), mouse monoclonal anti-22C10 (1:500; Developmental Studies Hybridoma Bank (DSHB)), mouse monoclonal anti-Armadillo (1:100; DSHB), and rabbit polyclonal anti-DPax2 (1:2000; J. Kavaler, Colby College). The secondary antibodies used were: Alexa Fluor® 488 goat anti-mouse and goat anti-rabbit (1:500; Cedarlane Labs), and TRITC goat anti-mouse (1:500; Cedarlane Labs). Staining embryos for f-actin using TRITC-phalloidin was performed as previously described (REED *et al*. 2001). Confocal microscopy and confocal image processing were performed as previously described (CORMIER *et al*. 2012). Preparation of embryos for live imaging was as previously described (REED *et al*. 2009).

### Fluorescent *in situ* hybridization (FISH)

Whole mount fluorescent *in situ* hybridization used 3 hour embryo collections of wild-type or *daGAL4* > *UAS-GFP-hnt* aged for 10 hours at 25° C, giving embryos at stage 13-16. Embryo fixation followed protocols as described (LECUYER *et al*. 2008). cDNA clones were acquired from the Drosophila Genomics Resource Center (Indiana University), including the *DPax2* clone IP01047.

### Cone cell distribution quantification

48hr APF pupal eye discs were immunostained using anti-armadillo as described above in three genetic backgrounds (*rl*, *peb*, *rl peb*). *peb* is a temperature sensitive recessive visible allele and was reared under permissive (25° C) and restrictive (29° C) conditions. *rl* and *rl peb* lines were reared at 25° C. Five to six independent eye discs were examined for each genotype and condition (*rl* 25° C, *peb* 25° C, *peb* 29° C, and *rl peb* 25° C). The average frequencies of cone cell within an ommatidium, ranging from 1-5, were calculated with the standard deviation then plotted onto a stacked bar graph.

### Recovery of *hnt^NP7278ex1^*

The viable and fertile *GAL4* enhancer trap line *NP7278*, inserted 158 bp upstream of the *hnt* transcription start site (THURMOND *et al*. 2019), was mobilized by crossing to *Δ2-3* transposase. Progeny were crossed to *FM7h, w B* and lines were established from single virgin females that had lost the *w^+^* marker of *NP7278*. Lethal lines (not producing *B^+^* progeny) were subsequently selected and tested for *GAL4* expression by crossing to *UAS-GFP^nls^*.

### *hnt^NP7278ex1^* rescue experiments

The *hnt^NP7278ex1^* stock was crossed into a background carrying second chromosome insertions *UAS-GFP^nls^* and *Ubi-DE-cadherin-GFP*. Virgin females of this resulting stock (*y w hnt^NP7278ex1^ FRT19A/ FM7h, w; UAS-GFP^nls^ Ubi-DE-cadherin-GFP/CyO*) were subsequently crossed to *tub-GAL80 hsFLP FRT19A* males (for control mutant) or to *tub-GAL80 hsFLP FRT19A; UAS-X* males for rescue experiments (where *UAS-*X was the homozygous *2^nd^* chromosome insertion *UAS-p35*, or one of the homozygous *3^rd^* chromosome insertions *UAS-sSpi*, *UAS-Ras85D^v12^*, or *UAS-pnt^P1^*). In the case of the *3^rd^* chromosome insertion *UAS-pnt^P2^*, which is not homozygous viable, male *tub-GAL80 hsFLP FRT19A; UAS-pnt^P2^ / UAS-Cherry^nls^* outcross progeny were used. Embryos between 12-14 hours old were collected from crosses of 30-40 females and males using an automated Drosophila egg collector (Flymax Scientific Ltd.) at room temperature (22°C) and mounted for live imaging as previously described (REED *et al*. 2009). For each imaging session, non-mutant embryos were confirmed as having completed or being in the terminal stages of dorsal closure. Mutant embryos (*hnt^NP7278ex1^/Y; UAS-GFP^nls^ Ubi-DE-cadherin-GFP/UAS-X* or *hnt^NP7278ex1^/Y; UAS-GFP^nls^ Ubi-DE-cadherin-GFP/+; UAS-X/+*) were unambiguously identified by expression of *UAS-GFP^nls^* (Fig. S3). In the case of *UAS-pnt^P2^*, mutant embryos also expressing *UAS-pnt^P2^* were identified as those embryos having *UAS-GFP^nls^* expression while lacking *UAS-Cherry^nls^* expression. A control rescue was performed by crossing to *y w hnt^XE81^ FRT19A; pebBAC^CH321-46J02^* males (BAC insert is *hnt^+^*). Images of mutant embryos were scored as one of three possible categories: 1) GBR failure (telson pointed anteriorly) with a small AS remnant; 2) GBR partial (telson pointed vertically or posteriorly but not at full posterior position) with an intact but distorted AS; 3) GBR complete (telson pointed posteriorly and located at normal posterior position) and with an intact but distorted or normal AS.

### Data and Reagent Availability

Stocks used that are unique to this study are available upon request. Supplemental material has been uploaded to figshare. The image data sets and embryo scoring result used to evaluate *hnt^NP7278ex1^* rescue (presented in Fig. 5K) are available as supplemental material (Fig. S1). Other supplemental material includes the demonstration of reduced *hnt* expression in *hnt^NP7278ex1^* mutant embryos (Fig. S2) and Punnett square diagrams detailing the genetic crosses used for the unambiguous identification of mutant and rescued *hnt^NP7278ex1^* mutant embryos (Fig. S3).

## Results

### PNS, chordotonal organ and oenocyte specification are disrupted in *hnt* loss-of-function mutants

In order to determine if phenotypes associated with reduced EGFR signaling are present in *hnt* mutants, we first examined the development of the PNS in *hnt^XE81^* mutant embryos using anti-Futsch/22C10 (hereafter referred to as 22C10), which labels all neurons of the PNS as well as some neurons of the central nervous system (CNS) (HUMMEL *et al*. 2000). *hnt^XE81^* mutant embryos lack sensory neurons (Fig. 1A, B). The absence of sensory neurons is most evident in the abdominal segments. Each embryonic abdominal hemisegment normally contains eight internal stretch receptors known as chordotonal organs, arranged as a single dorsal lateral organ (v’ch1), a lateral cluster of five (lch5), and two single ventral lateral organs (vchB, and vchA) (BREWSTER AND BODMER 1995). 22C10 immunostaining shows the neurons of the lch5 clusters are frequently reduced from five to three in number in *hnt^XE81^* mutants (asterisks, Fig. 1A, B and Fig. 1A’, B’). TRITC-phalloidin staining of f-actin confirms the reduction of the lch5 clusters from five to three (asterisks, Fig. 1C and Fig. 1D), and reveals a complete absence of the single chordotonal organs in *hnt^XE81^* mutants (arrowheads in Fig. 1C).

**Figure 1.**
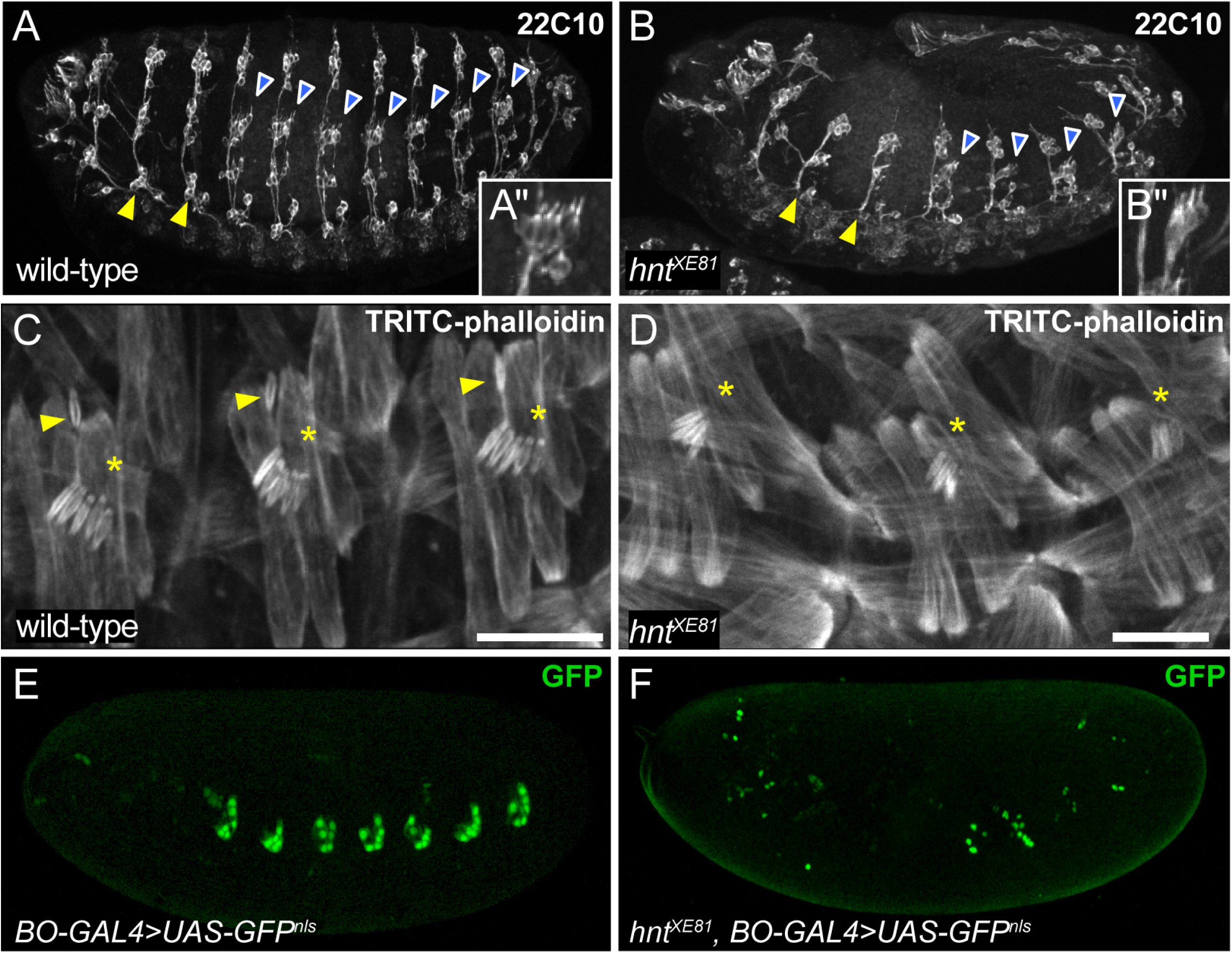
The embryonic *hnt* mutant phenotype includes hallmarks of reduced EGFR signaling. (**A**) Wild-type stage 15 embryo immunostained using the neuronal marker 22C10 showing typical development of the PNS, including clusters of ventral neurons in the second and third thoracic segments (arrowheads) and five neurons associated with lateral chordotonal organ clusters in the abdominal segments (blue with white outline arrowheads and inset A’). (**B**) 22C10 immunostained *hnt* mutant embryo showing the absence of neurons (arrowheads *cf.* panel A) including two of the five neurons of each lateral chordotonal cluster (blue with white outline arrowheads and inset B’). (**C**) TRITC-phalloidin stained stage 15 wild-type embryo showing the f-actin rich structure of the lateral chordotonal lch5 organ clusters (asterisks) and the dorsolateral chordotonal organ lch1 (arrowheads). (**D**) TRITC-phalloidin stained *hnt* mutant embryo showing differentiated lateral chordotonal organs that are reduced in number (asterisks) and the absence of the dorsolateral chordotonal lch1 organ. (**E**) Wild-type embryo showing *UAS-GFP^nls^* expression using the oenocyte-specific driver BO-GAL4. (**F**) *hnt^XE81^* mutant embryo showing reduced number of GFP-positive oenocytes (*BO-GAL4* > *UAS-GFP^nls^*) and failure to form oenocyte clusters. Scale bars represent 20 microns (C,D).

In general, mutants lacking lateral chordotonal organs do not form oenocytes, and EGFR signaling has been implicated in oenocyte induction (ELSTOB *et al*. 2001). We, therefore, used the oenocyte specific *BO-GAL4* to drive expression of *nuclear-GFP* in wild-type and *hnt^XE81^* mutants to evaluate oenocyte specification (Fig. 1E,F). In addition to *hnt* mutants having reduced numbers of *BO-GAL4*-positive cells, these cells are not organized into clusters as in wild-type, but are scattered throughout the mutant embryos. This newly reported phenotype of *hnt* mutants, that of missing chordotonal organs and a failure in oenocyte differentiation, is a hallmark of reduced EGFR signaling (MAKKI *et al*. 2014).

### *hnt^peb^* is enhanced by reduced MAPK

Given the above findings, we were next interested in determining if a genetic background of reduced EGFR signaling would enhance a *hnt* mutant phenotype. Using anti-Armadillo (Arm) immunostaining, we evaluated the pupal ommatidial structure of the temperature sensitive hypomorphic *hnt* allele *pebbled* (*hnt^peb^*) as well as a viable hypomorphic mutant of the EGFR downstream effector MAPK, also known as *rolled* (*rl^1^*). At the permissive temperature of 25°C, 87% of ommatidia in *hnt^peb^* mutants resemble wild-type and contain four cone cells (Fig. 2A,B *cf.* 2C; Fig. 2G). Likewise, 90% of ommatidia of *rl^1^* mutants raised at 25°C are normal (Fig. 2D,G). The number of ommatidia showing a normal cone cell number is reduced to 28% in *peb* mutants raised at the restrictive temperature of 29°C (Fig. 2E,G) while *peb; rl^1^* double mutants raised at the permissive temperature (25°C) display a distinct enhancement of the *peb* mutant phenotype, having only 22% of ommatidia with the correct cone cell number (Fig. 2F,G). These observations demonstrate a novel genetic interaction between *hnt* and *MAPK*, showing that *rl^1^* behaves as an enhancer of the cone cell specification defect of *hnt^peb^*. Interestingly, *hnt* is not expressed in cone cells, but is expressed in photoreceptor precursor cells (R cells) where it is required for induction and expression within cone cells of the determinant *DPax2* (PICKUP *et al*. 2009).

**Figure 2.**
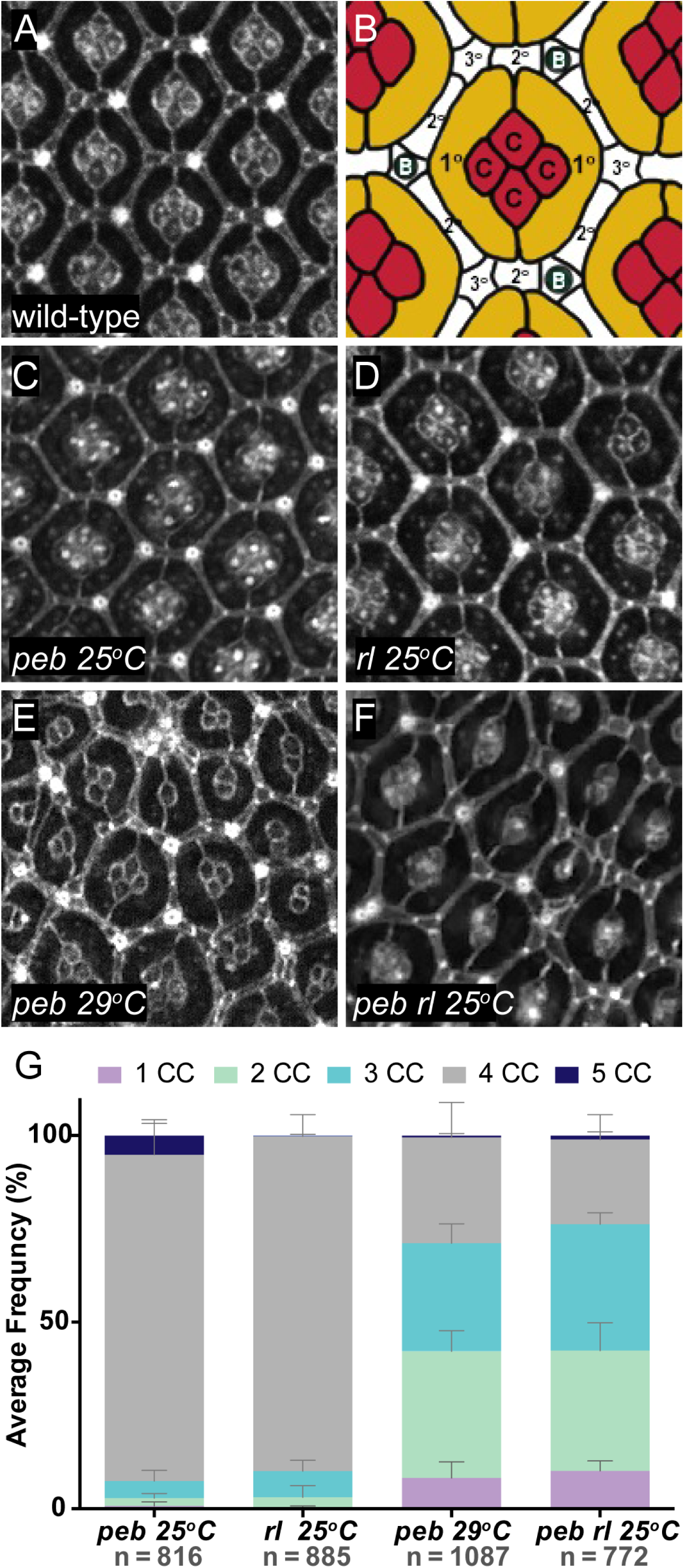
The viable temperature sensitive hypomorphic *hnt* allele *pebbled* (*hnt^peb^*) is enhanced by the viable hypomorphic MAPK allele *rolled* (*rl^1^*). (**A**) Anti-Arm immunostained wild-type pupal retina 48h APF showing the normal organization of ommatidial units. (**B**) Cartoon of wild-type ommatidial structure showing four cone cells (red - c), two primary pigment cells (yellow - 1°), and the secondary (white - 2°) and tertiary pigment cells (white - 3°) of the interommatidial lattice. Also depicted as a part of the lattice are the interommatidial bristles (dark green). (**C**) Anti-Arm immunostained pupal retina (48h APF) of *peb* mutant raised at the permissive temperature (25°C) showing normal ommatidial organization. (**D**) Anti-Arm immunostained pupal retina (48h APF) of *rl* mutant raised at 25°C showing normal ommatidial organization. (**E**) Anti-Arm immunostained pupal retina (48h APF) of *peb* mutant raised at the restrictive temperature (29°C) showing a disruption in ommatidial organization. (**F**) Anti-Arm immunostained pupal retina (48h APF) of *peb; rl* double mutant raised at the permissive temperature of 25°C showing disrupted ommatidial organization, indicating a genetic enhancement of *peb* under what is normally the permissive condition. (**G**) Stacked bar graph showing the average frequency of observed cone cells per ommatidium (1-5 CC) for *peb* 25°C, *rl* 25°C, *peb* 29°C, and *peb; rl* 25°C.

### Overexpression of *hnt* during embryogenesis results in ectopic *DPax2* expression

Using a candidate gene approach, we examined stage 13-16 embryos in which *UAS-GFP-hnt* was globally expressed using the *daGAL4* driver. Among candidate genes tested, *DPax2* (*CG11049*, also known as *shaven* (*sv*) or *sparkling* (*spa*)) was found to show a striking transcriptional upregulation in embryos overexpressing *hnt* compared to control embryos (Fig. 3A,B). The upregulation of *DPax2* in embryos overexpressing *hnt* was confirmed at the level of protein expression by anti-DPax2 immunostaining (Fig. 3C,D) as well as by reporter gene construct expression (Fig. 3E,F). Interestingly, *hnt* mutants do not abolish or reduce *DPax2* expression (Fig. 3G), suggesting that while *hnt* overexpression can result in *DPax2* overexpression, Hnt is not required for endogenous *DPax2* expression throughout the embryonic PNS.

**Figure 3.**
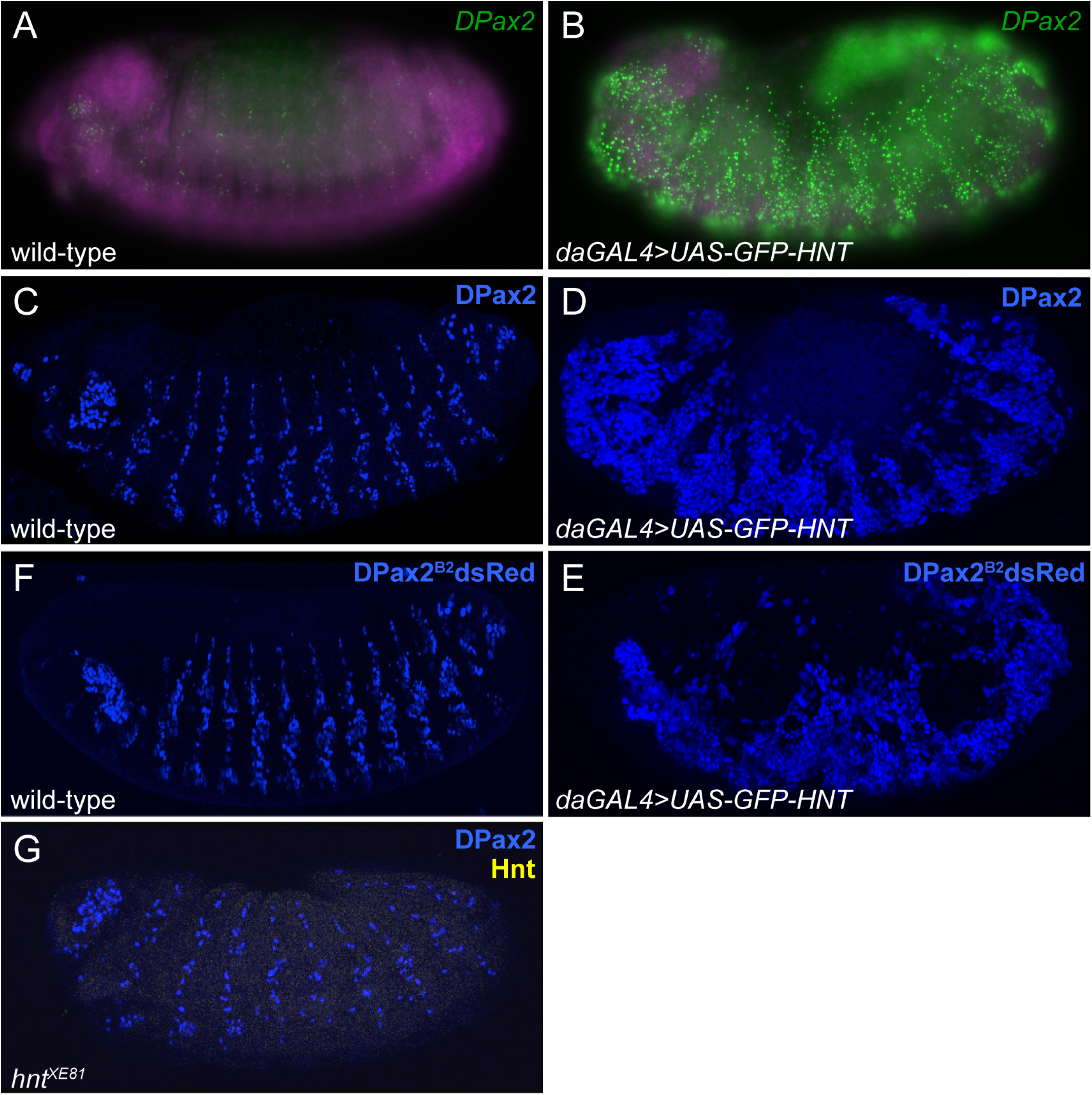
Global overexpression of *hnt* results in ectopic *DPax2* expression. (**A**) Wild-type embryo showing *DPax2* mRNA distribution expression using FISH (green) (**B**) Embryo overexpressing *hnt* (*daGAL4* > *UAS-GFP-hnt*) showing ectopic and increased levels of *DPax2* mRNA using FISH (green). (**C**) Wild-type embryo showing *DPax2* expression using anti-DPax2 immunostaining (blue). (**D**) Embryo overexpressing *hnt* immunostained for DPax2 (blue) showing ectopic DPax2 in large regions of lateral ectoderm. (**E**) Wild-type embryo showing expression of the *shaven* reporter gene construct *DPax2^B2^dsRed* (blue) as faithful to endogenous *DPax2* expression throughout the developing PNS. (**F**) Embryo overexpressing *hnt* showing ectopic *DPax2* expression using the *DPax2^B2^dsRed* reporter gene. (**G**) Embryo immunostained for DPax2 (blue) and Hnt (yellow) showing that this embryo is a *hnt^XE81^* mutant (absence of Hnt signal) and DPax2 throughout the PNS.

### Ectopic *DPax2* expression in the context of *hnt* overexpression is EGFR dependent

*DPax2* encodes a paired domain transcription factor and is expressed in the developing PNS, including the embryonic PNS, pupal eye, and micro- and macrochaetes (FU *et al*. 1998). We next wished to determine if *DPax2* expression in embryos overexpressing *hnt* is dependent on EGFR signaling. Compared to the overexpression control (Fig. 4A-A’’), we found that reduced EGFR (*Egfr^1a15^/Egfr^f2^*) suppresses ectopic *DPax2* expression (Fig. 4B-B’’). We also observed that *DPax2* overexpression associated with *hnt* overexpression is sensitive to *Egfr* dosage *as Egfr^f2^*/*+* heterozygous embryos show reduced *DPax2* expression relative to the overexpression control (Fig. 4C-C’’). To further corroborate *DPax2* ectopic expression as EGFR-dependent, we examined *DPax2* reporter gene expression in embryos globally expressing the activated EGFR ligand *secreted Spitz* (*sSpi*). Such embryos also show ectopic *DPax2* expression, suggesting that ectopic *DPax2* expression is elicited through increased EGFR signaling (Fig. 4 D,E). In addition, we found that the same *Egfr* mutant (*Egfr^1a15^/Egfr^f2^*) does show expression of the *DPax2^B2^dsRed* reporter. Although the total number of *DPax2* expressing cells is reduced relative to wildtype, this indicates that *Egfr* mutants are capable of producing cells that express *DPax2* (Fig. 4F). Taken together, these data are consistent with the interpretation that *DPax2* is not a direct target of *hnt*, that ectopic *DPax2* expression is a consequence of excessive EGFR signaling, and that *hnt* overexpression may result in *DPax2* overexpression through excessive EGFR signaling. Moreover, these results raise the possibility that *hnt* loss-of-function mutants could possibly be rescued by ectopic activation of Egfr signaling.

**Figure 4.**
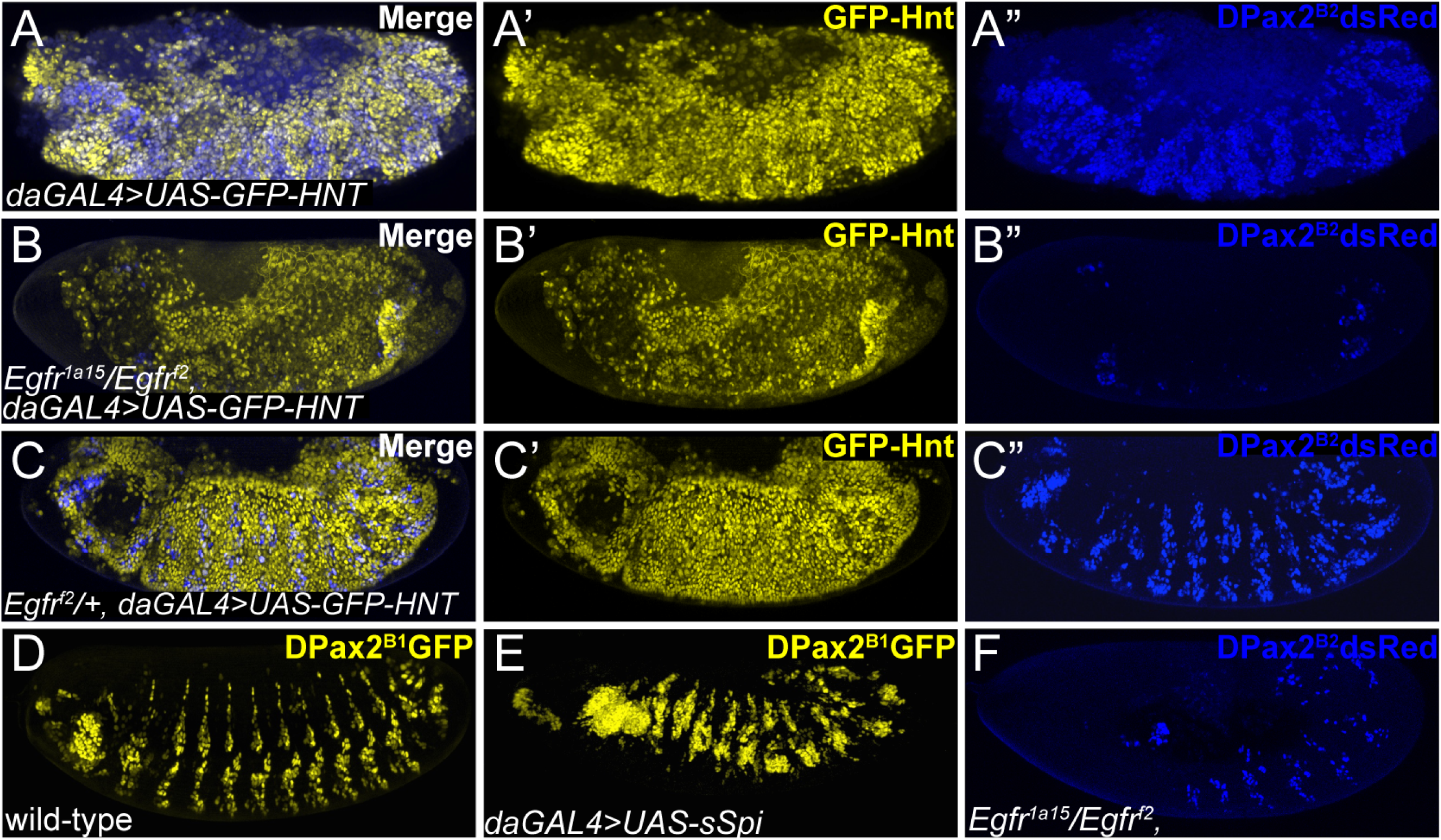
Ectopic *DPax2* expression associated with *hnt* overexpression requires EGFR signaling. (**A-A’’**) Immunostained *pan-GFP-hnt* embryo (*daGAL4* > *UAS-GFP-hnt*) showing Hnt (yellow, A’) and associated ectopic DPax2 (Blue, A’’). (**B-B’’**) *Pan-GFP-hnt* embryo that carries the loss-of-function allelic combination *Egfr^1a15^*/ *Egfr^f2^*, showing absence of ectopic *DPax2* expression using the *DPax2^B2^dsRed* reporter. (**C-C’’**) *Pan-GFP-hnt* embryo heterozygous for the *Egfr^f2^* allele showing reduced ectopic expression of the *DPax2^B2^dsRed* reporter. (**D**) Wild-type stage 15 embryo showing that expression of the *DPax2^B1^GFP* reporter gene is consistent with endogeneous DPax2 (*cf.* Fig. 3C). (**E**) Embryo expressing the *DPax2^B1^GFP* reporter gene in the background of globally activated EGFR signaling (*daGAL4* > *UAS-sSpi*) showing ectopic *DPax2* expression. (F) The loss-of-function allelic combination *Egfr^1a15^*/ *Egfr^f2^* in the absence of *hnt* overexpression, showing *DPax2* expression using the *DPax2^B2^dsRed* reporter.

### The embryonic U-shaped terminal mutant phenotype of *hnt^NP7278ex1^* is rescued by activation of EGFR signaling

Given the above results showing phenotypes related to reduced EGFR signaling in *hnt* mutants, the genetic enhancement between *hnt^peb^* and *rl^1^*, in addition to the EGFR-dependence of ectopic *DPax2* expression associated with *hnt* overexpression, we wished to test if *hnt* loss-of-function phenotypes can be rescued by activation of Egfr signaling. As is the case for *Egfr* mutants, *hnt* mutants fail to undergo or complete GBR and are associated with premature AS degeneration and death (FRANK AND RUSHLOW 1996; GOLDMAN-LEVI *et al*. 1996; LAMKA AND LIPSHITZ 1999). We conducted rescue experiments using a newly recovered *hnt* allele, *hnt^NP7278ex1^* (see Materials and Methods). The *hnt^NP7278ex1^* allele is a *GAL4* enhancer trap insertion that is embryonic lethal, fails to complement *hnt^XE81^*, shows premature AS degeneration, has GBR defects (Fig. 5D,E,K), and is rescued by *pebBAC^CH321-46J02^* (Fig. 5F, K). Very similar to the previously described allele *hnt^308^* (REED *et al*. 2001), *hnt^NP7278ex1^* shows reduced anti-Hnt immunostaining (Fig. S2). *hnt^NP7278ex1^* is, therefore, best characterized as a strong hypomorphic allele. Interestingly, the *hnt^NP7278ex1^* mutant retains *GAL4* expression in a pattern faithful to endogenous *hnt* expression, including early (prior to onset of GBR) expression in the AS (Fig 5A,B). The *hnt^NP7278ex1^* mutant phenotype, however, does not disrupt oenocyte specification or the lch5 cluster of chordotonal organs as we described for *hnt^XE81^*. We, therefore, chose to test for rescue of premature AS death and GBR failure. We were able to use *hnt^NP7278ex1^* in combination with an *X*-linked *tub-GAL80* insertion to unambiguously identify hemizygous *hnt^NP7278ex1^* mutant embryos that also express an autosomal UAS transgene (see Materials and Methods, and Fig. S3). We found that 72.4% (n=58) of control *hnt^NP7278ex1^* embryos show a strong U-shaped phenotype in which the AS is reduced to a small remnant, indicative of GBR failure and premature AS degeneration, respectively (Fig. 5E,K). The AS degeneration and GBR phenotype of *hnt^NP7278ex1^* mutants was rescued by expression of the baculovirus caspase inhibitor *UAS-p35* (5.9% GBR failure; n= 34; Fig. 5F,I), the activated EGFR ligand *UAS-sSpi* (0% GBR failure; n = 27, Fig. 5H,K), constitutively active RAS (8.3% GBR failure; n= 36; Fig. 5I,K). We also tested for rescue of *hnt^NP7278ex1^* by expression of two isoforms of the ETS transcription factor effector encoded by *pointed* (*pnt*), which is a downstream effector of the EGFR/Ras/MAPK pathway. The isoform Pnt^P2^ requires activation through phosphorylation by MAPK, whereas the Pnt^P1^ isoform, which is transcriptionally activated by the activated form of Pnt^P2^, is constitutively active without activation by MAPK (O’NEILL *et al*. 1994; SHWARTZ *et al*. 2013). Expression of the constitutively active isoform via *UAS*-*Pnt^P1^* resulted in rescue (9.1% GBR failure; n= 31; Fig.5J,K). Interestingly, expression the other isoform via *UAS-Pnt^P2^* did not rescue *hnt^NP7278ex^* (72.0% GBR failure, n= 25; Fig. 5K). All image data sets and scoring annotations used to generate Fig. 5K are presented as supplemental material (Fig. S1). Rescue by *UAS-p35* confirms that premature AS degeneration in *hnt* mutants is associated with caspase activation. Furthermore, rescue of *hnt* mutants by expression of components of the EGFR signaling pathway is consistent with *hnt* operating either upstream or in parallel to this pathway. Rescue was not complete in that AS morphology was abnormal, and rescued embryos failed to complete dorsal closure likely due to the abnormal persistence of the rescued AS. Interestingly, the failure to rescue AS death and GBR defects by expression of the *Pnt^P2^* isoform, which requires activation through phosphorylation by MAPK (O’NEILL *et al*. 1994; SHWARTZ *et al*. 2013), is consistent with reduced MAPK activity within the AS of *hnt* mutants.

**Figure 5.**
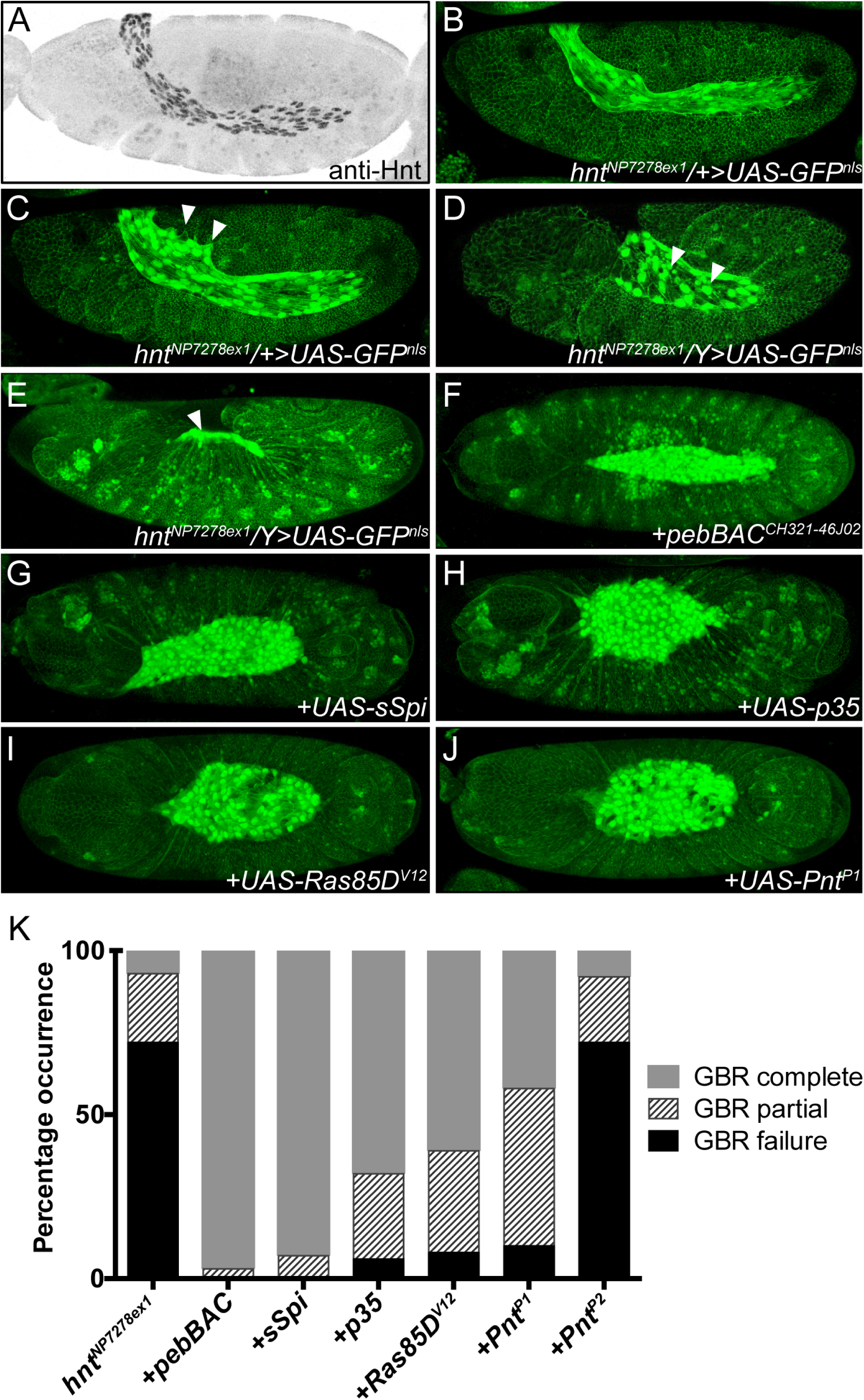
GBR and premature amnioserosa death of *hnt^NP7278ex1^* is rescued by caspase suppression and by activation of EGFR signaling. (**A**) Anti-Hnt immunostained showing AS expression prior to onset of GBR. (**B**) Live confocal image of *hnt^NP7278ex1^*/*+*; *UAS-GFP^nls^ Ubi-DEcadherin-GFP*/*+* embryo showing AS expression associated with *hnt^NP7278ex1^* prior to onset of GBR. (**C**) Same embryo shown in B imaged 67 minutes later during initiation of GBR. The AS is folded over the extended tail and lamellopodia-type extensions contact the epidermis (white arrowheads. (**D**) Live confocal image of *hnt^NP7278ex1^*/*Y*; *UAS-GFP^nls^ Ubi-DEcadherin-GFP*/*+* mutant embryo at onset of GBR showing a failure of AS to maintain the fold over the posterior tail. AS apoptotic corpses are also present (white arrowheads). (**E**) Terminal GBR failure phenotype of *hnt^NP7278ex1^*/Y; *UAS-GFP^nls^ Ubi-DEcadherin-GFP*/+ mutant embryo showing tail-up phenotype and AS remnant (white arrowhead). (**F**) Control rescue embryo: *hnt^NP7278ex1^* or *hnt^NP7278ex1^*/*hnt^XE81^* mutant with *UAS-GFPnls Ubi-DEcadherin* showing rescue by *pebBAC^CH321-46J02^*. (**G**) GBR complete rescue of *hnt^NP7278ex1^* by *UAS-sSpi*. (**H**) GBR complete rescue of *hnt^NP7278ex1^* by *UAS-p35*. (**I**) GBR complete rescue of *hnt^NP7278ex1^* by *UAS-Ras85D^V12^*. (**J**) GBR complete rescue of *hnt^NP7278ex1^ by UAS-pnt^P1^*. (**K**) Stacked bar graph showing the frequency of GBR defects in *hnt^NP7278ex1^* mutants and rescue backgrounds.

## Discussion

### *hnt* loss-of-function and *hnt* overexpression phenotypes are consistent with perturbations in EGFR signaling

The development of chordotonal organs and oenocyte specification are both disrupted in *hnt* mutants and these phenotypes are hallmarks of reduced EGFR signaling. As an overview, each embryonic abdominal hemisegment normally develops eight chordotonal organs, organized into three single organs (v’ch1, vchB, and vchA), and a cluster of five organs (lch5). The embryonic specification and differentiation of chordotonal organs initiates with the delamination of chordotonal precursor cells (COPs) from the ectoderm (reviewed in (GOULD *et al*. 2001)). Briefly, chordotonal organs arise from five primary COPs (C1-C5), where C1-C3 give rise to the five organs of lch5, C4 is a precursor of v’ch1, and C5 is the precursor for vchB and vchA. The secretion of the active EGFR ligand Spitz by C3 and C5 expands the number of COPs from five to eight. Further EGFR signaling elicited by the C1 COP is also required for the induction of oenocytes (reviewed in (MAKKI *et al*. 2014)). In the absence of Egfr signaling, C1 fails to recruit oenocytes, and C3 fails to recruit secondary COPs to complete the five lateral chordotonal organs of the lch5 cluster (GOULD *et al*. 2001). Mutant phenotypes of genes belonging to what has been called the Spitz group (which encode components of the EGFR signaling pathway and include *Star*, *rhomboid*, *spitz*, and *pointed*), as well as the expression of dominant-negative EGFR, all display an absence of oenocytes and the formation of only three lateral chordotonal organs within the lch5 cluster (BIER *et al*. 1990; ELSTOB *et al*. 2001; RUSTEN *et al*. 2001). Based on our analysis of *hnt* mutant embryos, we suggest that *hnt* can be aptly described as a previously unrecognized member of the Spitz group of mutants. Overall, however, our findings represent additions to the list of phenotypic similarities between *hnt* and *Egfr* mutants, including germ band retraction and dorsal closure failure, as well as the loss of tracheal epithelial integrity (CLIFFORD AND SCHUPBACH 1992; CELA AND LLIMARGAS 2006; SHEN *et al*. 2013).

We found *hnt* overexpression in the embryo results in increased and ectopic expression of *DPax2*, and we found this effect to be unequivocally Egfr-dependent. We also found that global activation of Egfr signaling via expression of the Egfr ligand *sSpi* also causes *DPax2* overexpression. Our results are consistent with previous work showing that Hnt is required in the developing eye imaginal disc for cone cell induction; here, it was also shown that reduced *hnt* expression resulted in reduced DPax2, that *hnt* overexpression resulted in increased DPax2, and that these effects were non-autonomous (PICKUP *et al*. 2009). The suggested model was that Hnt is required within the R1/R6 photoreceptor precursor cells to achieve a level of Delta sufficient for cone cell induction. While our suggestion that Hnt promotes Egfr signaling is not mutually exclusive with a role in promoting *Delta* expression, it is noteworthy that the expression of *Delta* within R-precursor cells is elevated by the activation of EGFR signaling in these cells (TSUDA *et al*. 2006). The observation of reduced Delta associated with reduced *hnt* expression could, therefore, be attributed to reduced Hnt-dependent EGFR signaling within the R-precursor cells.

### Rescue of the *hnt* U-shaped mutant phenotype

The AS, which is programmed to die during and following the process of dorsal closure, is possibly required for mechanical as well as signaling events that are critical for the morphogenetic processes of GBR and dorsal closure. Premature AS death may, therefore, lead to U-shaped or dorsal closure phenotypes. In support of this view, AS-specific cell abalation disrupts dorsal closure (SCUDERI AND LETSOU 2005), and other U-shaped mutants display premature AS death, including *u-shaped* (*ush*), *tail-up* (*tup*), *serpent* (*srp*), and *myospheroid* (*mys*) (FRANK AND RUSHLOW 1996; GOLDMAN-LEVI *et al*. 1996; REED *et al*. 2004).

AS programmed cell death normally occurs through an upregulation of autophagy in combination with caspase activation (MOHSENI *et al*. 2009; CORMIER *et al*. 2012). AS death can be prevented, resulting in a persistent AS phenotype, in a number of backgrounds. These include expression of the caspase inhibitor *p35*, RNAi knockdown of the proapoptotic gene *hid*, expression of activated Insulin receptor (*dInR^ACT^*), dominant negative ecdysone receptor (*EcR^DN^*), active EGFR ligand *secreted Spitz* (*sSpi*), constitutively active RAS (*Ras85D^V12^*), as well as over expression of *Egfr-GFP* (MOHSENI *et al*. 2009; SHEN *et al*. 2013). In addition, embryos homozygous for *Df(3L)H99*, which deletes the pro-apoptotic gene cluster *reaper/hid/grim*, also present a persistent AS phenotype (MOHSENI *et al*. 2009; CORMIER *et al*. 2012). During normal development, Hnt is no longer detectable by immunostaining within the AS as it begins to degenerate following dorsal closure (REED *et al*. 2004; MOHSENI *et al*. 2009). Thus, it is likely that *hnt* downregulation is required for normal AS degeneration, and that the mutant phenotype of *hnt* is the result of a premature activation of the normal death process. In support of this, we have demonstrated that several backgrounds associated with a persistent AS phenotype are able to rescue GBR failure and AS death in *hnt* mutants.

In the context of programmed cell death within the embryonic CNS, MAPK dependent phosphorylation has been show to inhibit the pro-apoptotic activity of the Hid protein (BERGMANN *et al*. 2002). We suggest that Egfr signaling within the AS could also represent a survival signal, leading to MAPK activation and Hid inhibition. Several observations are consistent with this model, including AS expression of several components of the Egfr signaling pathway. For example, within the AS anlage there is robust expression of *rhomboid* (*rho*) (FRANCOIS *et al*. 1994), which encodes a intramembrane serine protease required for the activation of EGFR ligands; see (SHILO 2005). In addition, prior to the onset of GBR, there is pronounced AS expression of *vein* (*vn*), which encodes an additional EGFR ligand (SCHNEPP *et al*. 1996). Vein is a weaker EGFR ligand, but it is produced in an active form and is not subject to inhibition by the EGFR antagonist Argos (Aos); see (GOLEMBO *et al*. 1999; SHILO 2005). At about the same stage, expression of a downstream EGFR effector *pointed* (*pnt*) is found in the AS, as is *hid,* which is also expressed in the apoptotic AS (see Berkeley Drosophila Genome Project; https://insitu.fruitfly.org/cgi-bin/ex/insitu.pl).

### Potential Hnt target genes and EGFR signaling

As a model for normal AS death, we suggest that a downregulation of *hnt* expression could lead to reduced EGFR AS signaling, thereby decreasing MAPK inhibitory phosphorylation of the pro-apoptotic protein Hid. According to this model, AS death and subsequent GBR failure in *hnt* mutants would be attributed to reduced EGFR signaling, lower MAPK activity, and pro-apoptotic activity of unphosphorylated Hid.

But how might *hnt* expression promote Egfr signaling and maintain high MAPK activity? A recent genetic screen for genes involved in the regulation of Wallerian degeneration (the fragmentation and clearance of severed axons) identified *hnt* as being required for this process. As part of this work, the authors performed ChIP-seq analysis of a *GM2* Drosophila cell line expressing a tagged version of Hnt. This resulted in the identification of 80 potential direct targets of Hnt (FARLEY *et al*. 2018). Interestingly, several of these putative Hnt target genes are also known targets of the EGFR signaling pathway, including *InR* (ZHANG *et al*. 2011), *E2f1* (XIANG *et al*. 2017), *bantam* (HERRANZ *et al*. 2012), *Dl* (TSUDA *et al*. 2002), and *dve* (SHIRAI *et al*. 2003); while others have been implicated in the regulation of EGFR signaling and include *EcR* (QIAN *et al*. 2014), *srp* (CAMPBELL *et al*. 2018), *MESR6* (HUANG AND RUBIN 2000), *Madm* (SINGH *et al*. 2016), and *skd* (LIM *et al*. 2007). Also, and of particular interest, among the genes identified are known target genes of EGFR signaling that are also regulators or effectors of EGFR signaling. These include the gene *pnt*, which encodes an ETS transcriptional activator - a key component for the transcriptional output of EGFR signaling that can also create a positive feedback loop through the transcription of *vn* (GOLEMBO *et al*. 1999; PAUL *et al*. 2013; CRUZ *et al*. 2015), and *Mkp3* (*Mitogen-activated protein kinase*), which is a negative regulator of EGFR signaling (GABAY *et al*. 1996; KIM *et al*. 2004; BUTCHAR *et al*. 2012). Further investigations will be required to determine if the phenotypes associated with *hnt* overexpression, as well as *hnt* loss-of-function, can be attributable (in whole or in part) to changes in expression of any of these potential target genes.

## Acknowledgements

We thank the Bloomington Drosophila Resource Center (BDRC) and the Kyoto Drosophila Resource Center for genetic stocks. We are grateful to M. Freeman, A. Gould, N. Harden, A. Hilliker, and H. Lipshitz for additional stocks and reagents. We also thank the Developmental Studies Hybridoma Bank (DSHB), created by the NICHD of the NIH and maintained at The University of Iowa, Department of Biology, Iowa City, IA 52242. We thank the Drosophila Genomics Resource Center (Indiana University). H.M.K. was supported by the Canadian Institute of Health Research (MOP 133473) and B.H.R. was supported by the Natural Sciences and Engineering Research Council of Canada (NSERC RGPIN-2015-04458).

## References

1. Bachmann, A., and E. Knust, 2008 The use of P-element transposons to generate transgenic flies. Methods Mol Biol 420: 61–77.

2. Baechler, B. L., C. McKnight, P. C. Pruchnicki, N. A. Biro and B. H. Reed, 2015 Hindsight/RREB-1 functions in both the specification and differentiation of stem cells in the adult midgut of Drosophila. Biology Open.

3. Barolo, S., B. Castro and J. W. Posakony, 2004 New Drosophila transgenic reporters: insulated P-element vectors expressing fast-maturing RFP. Biotechniques 36: 436–440, 442.

4. Bergmann, A., M. Tugentman, B. Z. Shilo and H. Steller, 2002 Regulation of cell number by MAPK-dependent control of apoptosis: a mechanism for trophic survival signaling. Dev Cell 2: 159–170.

5. Bier, E., L. Y. Jan and Y. N. Jan, 1990 rhomboid, a gene required for dorsoventral axis establishment and peripheral nervous system development in Drosophila melanogaster. Genes Dev 4: 190–203.

6. Brewster, R., and R. Bodmer, 1995 Origin and specification of type II sensory neurons in Drosophila. Development 121: 2923–2936.

7. Brodu, V., P. R. Elstob and A. P. Gould, 2004 EGF receptor signaling regulates pulses of cell delamination from the Drosophila ectoderm. Dev Cell 7: 885–895.

8. Buffin, E., and M. Gho, 2010 Laser microdissection of sensory organ precursor cells of Drosophila microchaetes. PLoS One 5: e9285.

9. Butchar, J. P., D. Cain, S. N. Manivannan, A. D. McCue, L. Bonanno et al., 2012 New negative feedback regulators of Egfr signaling in Drosophila. Genetics 191: 1213–1226.

10. Campbell, K., G. Lebreton, X. Franch-Marro and J. Casanova, 2018 Differential roles of the Drosophila EMT-inducing transcription factors Snail and Serpent in driving primary tumour growth. PLoS Genet 14: e1007167.

11. Cela, C., and M. Llimargas, 2006 Egfr is essential for maintaining epithelial integrity during tracheal remodelling in Drosophila. Development 133: 3115–3125.

12. Clifford, R., and T. Schupbach, 1992 The torpedo (DER) receptor tyrosine kinase is required at multiple times during Drosophila embryogenesis. Development 115: 853–872.

13. Cormier, O., N. Mohseni, I. Voytyuk and B. H. Reed, 2012 Autophagy can promote but is not required for epithelial cell extrusion in the amnioserosa of the Drosophila embryo. Autophagy 8: 252–264.

14. Cruz, J., N. Bota-Rabassedas and X. Franch-Marro, 2015 FGF coordinates air sac development by activation of the EGF ligand Vein through the transcription factor PntP2. Sci Rep 5: 17806.

15. Deady, L. D., W. Li and J. Sun, 2017 The zinc-finger transcription factor Hindsight regulates ovulation competency of Drosophila follicles. Elife 6.

16. Elstob, P. R., V. Brodu and A. P. Gould, 2001 spalt-dependent switching between two cell fates that are induced by the Drosophila EGF receptor. Development 128: 723–732.

17. Farley, J. E., T. C. Burdett, R. Barria, L. J. Neukomm, K. P. Kenna et al., 2018 Transcription factor Pebbled/RREB1 regulates injury-induced axon degeneration. Proc Natl Acad Sci U S A 115: 1358–1363.

18. Francois, V., M. Solloway, J. W. O’Neill, J. Emery and E. Bier, 1994 Dorsal-ventral patterning of the Drosophila embryo depends on a putative negative growth factor encoded by the short gastrulation gene. Genes Dev 8: 2602–2616.

19. Frank, L. H., and C. Rushlow, 1996 A group of genes required for maintenance of the amnioserosa tissue in Drosophila. Development 122: 1343–1352.

20. Franklin, R. B., J. Zou and L. C. Costello, 2014 The cytotoxic role of RREB1, ZIP3 zinc transporter, and zinc in human pancreatic adenocarcinoma. Cancer Biol Ther 15: 1431-1437.

21. Fu, W., H. Duan, E. Frei and M. Noll, 1998 shaven and sparkling are mutations in separate enhancers of the Drosophila Pax2 homolog. Development 125: 2943–2950.

22. Gabay, L., H. Scholz, M. Golembo, A. Klaes, B. Z. Shilo et al., 1996 EGF receptor signaling induces pointed P1 transcription and inactivates Yan protein in the Drosophila embryonic ventral ectoderm. Development 122: 3355–3362.

23. Goldman-Levi, R., C. Miller, G. Greenberg, E. Gabai and N. B. Zak, 1996 Cellular pathways acting along the germband and in the amnioserosa may participate in germband retraction of the Drosophila melanogaster embryo. Int J Dev Biol 40: 1043–1051.

24. Golembo, M., T. Yarnitzky, T. Volk and B. Z. Shilo, 1999 Vein expression is induced by the EGF receptor pathway to provide a positive feedback loop in patterning the Drosophila embryonic ventral ectoderm. Genes Dev 13: 158–162.

25. Gould, A. P., P. R. Elstob and V. Brodu, 2001 Insect oenocytes: a model system for studying cell-fate specification by Hox genes. J Anat 199: 25–33.

26. Gutierrez, E., D. Wiggins, B. Fielding and A. P. Gould, 2007 Specialized hepatocyte-like cells regulate Drosophila lipid metabolism. Nature 445: 275–280.

27. Herranz, H., X. Hong and S. M. Cohen, 2012 Mutual repression by bantam miRNA and Capicua links the EGFR/MAPK and Hippo pathways in growth control. Curr Biol 22: 651–657.

28. Huang, A. M., and G. M. Rubin, 2000 A misexpression screen identifies genes that can modulate RAS1 pathway signaling in Drosophila melanogaster. Genetics 156: 1219–1230.

29. Hummel, T., K. Krukkert, J. Roos, G. Davis and C. Klambt, 2000 Drosophila Futsch/22C10 is a MAP1B-like protein required for dendritic and axonal development. Neuron 26: 357–370.

30. Johnson, S. A., K. J. Harmon, S. G. Smiley, F. M. Still and J. Kavaler, 2011 Discrete regulatory regions control early and late expression of D-Pax2 during external sensory organ development. Dev Dyn 240: 1769–1778.

31. Kent, O. A., K. Fox-Talbot and M. K. Halushka, 2013 RREB1 repressed miR-143/145 modulates KRAS signaling through downregulation of multiple targets. Oncogene 32: 2576–2585.

32. Kent, O. A., M. N. McCall, T. C. Cornish and M. K. Halushka, 2014 Lessons from miR-143/145: the importance of cell-type localization of miRNAs. Nucleic Acids Res 42: 7528–7538.

33. Kim, M., G. H. Cha, S. Kim, J. H. Lee, J. Park et al., 2004 MKP-3 has essential roles as a negative regulator of the Ras/mitogen-activated protein kinase pathway during Drosophila development. Mol Cell Biol 24: 573–583.

34. Krejci, A., F. Bernard, B. E. Housden, S. Collins and S. J. Bray, 2009 Direct response to Notch activation: signaling crosstalk and incoherent logic. Sci Signal 2: ra1.

35. Lamka, M. L., and H. D. Lipshitz, 1999 Role of the amnioserosa in germ band retraction of the Drosophila melanogaster embryo. Dev Biol 214: 102–112.

36. Lecuyer, E., A. S. Necakov, L. Caceres and H. M. Krause, 2008 High-resolution fluorescent in situ hybridization of Drosophila embryos and tissues. CSH Protoc 2008: pdb prot5019.

37. Lim, J., O. K. Lee, Y. C. Hsu, A. Singh and K. W. Choi, 2007 Drosophila TRAP230/240 are essential coactivators for Atonal in retinal neurogenesis. Dev Biol 308: 322–330.

38. Liu, H., H. C. Hew, Z. G. Lu, T. Yamaguchi, Y. Miki et al., 2009 DNA damage signalling recruits RREB-1 to the p53 tumour suppressor promoter. Biochem J 422: 543–551.

39. Makki, R., E. Cinnamon and A. P. Gould, 2014 The development and functions of oenocytes. Annu Rev Entomol 59: 405–425.

40. Melani, M., K. J. Simpson, J. S. Brugge and D. Montell, 2008 Regulation of cell adhesion and collective cell migration by hindsight and its human homolog RREB1. Curr Biol 18: 532–537.

41. Ming, L., R. Wilk, B. H. Reed and H. D. Lipshitz, 2013 Drosophila Hindsight and mammalian RREB-1 are evolutionarily conserved DNA-binding transcriptional attenuators. Differentiation 86: 159–170.

42. Mohseni, N., S. C. McMillan, R. Chaudhary, J. Mok and B. H. Reed, 2009 Autophagy promotes caspase-dependent cell death during Drosophila development. Autophagy 5: 329–338.

43. Mukhopadhyay, N. K., B. Cinar, L. Mukhopadhyay, M. Lutchman, A. S. Ferdinand et al., 2007 The zinc finger protein ras-responsive element binding protein-1 is a coregulator of the androgen receptor: implications for the role of the Ras pathway in enhancing androgenic signaling in prostate cancer. Mol Endocrinol 21: 2056–2070.

44. O’Neill, E. M., I. Rebay, R. Tjian and G. M. Rubin, 1994 The activities of two Ets-related transcription factors required for Drosophila eye development are modulated by the Ras/MAPK pathway. Cell 78: 137–147.

45. Oliva, C., C. Molina-Fernandez, M. Maureira, N. Candia, E. Lopez et al., 2015 Hindsight regulates photoreceptor axon targeting through transcriptional control of jitterbug/Filamin and multiple genes involved in axon guidance in Drosophila. Dev Neurobiol.

46. Paul, L., S. H. Wang, S. N. Manivannan, L. Bonanno, S. Lewis et al., 2013 Dpp-induced Egfr signaling triggers postembryonic wing development in Drosophila. Proc Natl Acad Sci U S A 110: 5058–5063.

47. Pickup, A. T., M. L. Lamka, Q. Sun, M. L. Yip and H. D. Lipshitz, 2002 Control of photoreceptor cell morphology, planar polarity and epithelial integrity during Drosophila eye development. Development 129: 2247–2258.

48. Pickup, A. T., L. Ming and H. D. Lipshitz, 2009 Hindsight modulates Delta expression during Drosophila cone cell induction. Development 136: 975–982.

49. Pitsouli, C., and N. Perrimon, 2010 Embryonic multipotent progenitors remodel the Drosophila airways during metamorphosis. Development 137: 3615–3624.

50. Qian, Y., N. Dominado, R. Zoller, C. Ng, K. Kudyba et al., 2014 Ecdysone signaling opposes epidermal growth factor signaling in regulating cyst differentiation in the male gonad of Drosophila melanogaster. Dev Biol 394: 217–227.

51. Reed, B. H., S. C. McMillan and R. Chaudhary, 2009 The preparation of Drosophila embryos for live-imaging using the hanging drop protocol. J Vis Exp.

52. Reed, B. H., R. Wilk and H. D. Lipshitz, 2001 Downregulation of Jun kinase signaling in the amnioserosa is essential for dorsal closure of the Drosophila embryo. Curr Biol 11: 1098–1108.

53. Reed, B. H., R. Wilk, F. Schock and H. D. Lipshitz, 2004 Integrin-dependent apposition of Drosophila extraembryonic membranes promotes morphogenesis and prevents anoikis. Curr Biol 14: 372–380.

54. Reeves, N., and J. W. Posakony, 2005 Genetic programs activated by proneural proteins in the developing Drosophila PNS. Dev Cell 8: 413–425.

55. Rusten, T. E., R. Cantera, J. Urban, G. Technau, F. C. Kafatos et al., 2001 Spalt modifies EGFR-mediated induction of chordotonal precursors in the embryonic PNS of Drosophila promoting the development of oenocytes. Development 128: 711–722.

56. Schnepp, B., G. Grumbling, T. Donaldson and A. Simcox, 1996 Vein is a novel component in the Drosophila epidermal growth factor receptor pathway with similarity to the neuregulins. Genes Dev 10: 2302–2313.

57. Scuderi, A., and A. Letsou, 2005 Amnioserosa is required for dorsal closure in Drosophila. Dev Dyn 232: 791–800.

58. Shen, W., X. Chen, O. Cormier, D. C. Cheng, B. Reed et al., 2013 Modulation of morphogenesis by Egfr during dorsal closure in Drosophila. PLoS One 8: e60180.

59. Shen, W., and J. Sun, 2017 Dynamic Notch Signaling Specifies Each Cell Fate in Drosophila Spermathecal Lineage. G3 (Bethesda) 7: 1417-1427.

60. Shilo, B. Z., 2005 Regulating the dynamics of EGF receptor signaling in space and time. Development 132: 4017–4027.

61. Shirai, T., A. Maehara, N. Kiritooshi, F. Matsuzaki, H. Handa et al., 2003 Differential requirement of EGFR signaling for the expression of defective proventriculus gene in the Drosophila endoderm and ectoderm. Biochem Biophys Res Commun 311: 473–477.

62. Shwartz, A., S. Yogev, E. D. Schejter and B. Z. Shilo, 2013 Sequential activation of ETS proteins provides a sustained transcriptional response to EGFR signaling. Development 140: 2746–2754.

63. Singh, S. R., Y. Liu, J. Zhao, X. Zeng and S. X. Hou, 2016 The novel tumour suppressor Madm regulates stem cell competition in the Drosophila testis. Nat Commun 7: 10473.

64. Sun, J., and W. M. Deng, 2007 Hindsight mediates the role of notch in suppressing hedgehog signaling and cell proliferation. Dev Cell 12: 431-442.

65. Terriente-Felix, A., J. Li, S. Collins, A. Mulligan, I. Reekie et al., 2013 Notch cooperates with Lozenge/Runx to lock haemocytes into a differentiation programme. Development 140: 926–937.

66. Thiagalingam, A., A. De Bustros, M. Borges, R. Jasti, D. Compton et al., 1996 RREB-1, a novel zinc finger protein, is involved in the differentiation response to Ras in human medullary thyroid carcinomas. Mol Cell Biol 16: 5335–5345.

67. Thurmond, J., J. L. Goodman, V. B. Strelets, H. Attrill, L. S. Gramates et al., 2019 FlyBase 2.0: the next generation. Nucleic Acids Res 47: D759–D765.

68. Tsuda, L., M. Kaido, Y. M. Lim, K. Kato, T. Aigaki et al., 2006 An NRSF/REST-like repressor downstream of Ebi/SMRTER/Su(H) regulates eye development in Drosophila. EMBO J 25: 3191–3202.

69. Tsuda, L., R. Nagaraj, S. L. Zipursky and U. Banerjee, 2002 An EGFR/Ebi/Sno pathway promotes delta expression by inactivating Su(H)/SMRTER repression during inductive notch signaling. Cell 110: 625–637.

70. Wieschaus, E., C. Nüsslein-Volhard and G. Jürgens, 1984 Mutations affecting the pattern of the larval cuticle inDrosophila melanogaster. Wilhelm Roux’s Archives of Developmental Biology 193: 296–307.

71. Wilk, R., A. T. Pickup, J. K. Hamilton, B. H. Reed and H. D. Lipshitz, 2004 Dose-sensitive autosomal modifiers identify candidate genes for tissue autonomous and tissue nonautonomous regulation by the Drosophila nuclear zinc-finger protein, hindsight. Genetics 168: 281–300.

72. Wilk, R., B. H. Reed, U. Tepass and H. D. Lipshitz, 2000 The hindsight gene is required for epithelial maintenance and differentiation of the tracheal system in Drosophila. Dev Biol 219: 183–196.

73. Xiang, J., J. Bandura, P. Zhang, Y. Jin, H. Reuter et al., 2017 EGFR-dependent TOR-independent endocycles support Drosophila gut epithelial regeneration. Nat Commun 8: 15125.

74. Yip, M. L., M. L. Lamka and H. D. Lipshitz, 1997 Control of germ-band retraction in Drosophila by the zinc-finger protein HINDSIGHT. Development 124: 2129–2141.

75. Zhang, W., B. J. Thompson, V. Hietakangas and S. M. Cohen, 2011 MAPK/ERK signaling regulates insulin sensitivity to control glucose metabolism in Drosophila. PLoS Genet 7: e1002429.

